# Batch-Mask: An automated Mask R-CNN workflow to isolate non-standard biological specimens for color pattern analysis

**DOI:** 10.1101/2021.11.12.468394

**Authors:** John David Curlis, Timothy Renney, Alison R. Davis Rabosky, Talia Y. Moore

## Abstract

1. Efficient comparisons of biological color patterns are critical for understanding the mechanisms by which organisms evolve in ecosystems, including sexual selection, predator-prey interactions, and thermoregulation. However, elongate or spiral-shaped organisms do not conform to the standard orientation and photographic techniques required for automated analysis. Currently, large-scale color analysis of elongate animals requires time-consuming manual landmarking, which reduces their representation in coloration research despite their ecological importance.
2. We present `Batch-Mask`: an automated and customizable workflow to facilitate the analysis of large photographic data sets of non-standard biological subjects. First, we present a user guide to run an open-source region-based convolutional neural network with fine-tuned weights for identifying and isolating a biological subject from a background (masking). Then, we demonstrate how to combine masking with existing manual visual analysis tools into a single streamlined, automated workflow for comparing color patterns across images.
3. `Batch-Mask` was 60x faster than manual landmarking, produced masks that correctly identified 96% of all snake pixels, and produced pattern energy results that were not significantly different from the manually landmarked data set.
4. The fine-tuned weights for the masking neural network, user guide, and automated workflow substantially decrease the amount of time and attention required to quantitatively analyze non-standard biological subjects. By using these tools, biologists will be able to compare color, pattern, and shape differences in large data sets that include significant morphological variation in elongate body forms. This advance will be especially valuable for comparative analyses of natural history collections, and through automation can greatly expand the scale of space, time, or taxonomic breadth across which color variation can be quantitatively examined.

## 1 INTRODUCTION

The increasing digitization of museum specimens and the convenience of digital photography provide unparalleled opportunity to quantify how color varies across the entire tree of life. However, high morphological shape variation across taxa poses a challenge for automated image analysis tools, requiring prohibitively labor-intensive analysis with manual approaches. Snakes in particular demonstrate impressive variation in coloration and patterning (Allen et al., 2013) that serve critical functions, such as anti-predator signaling (Brodie III, 1993). Despite the iconic role of snake coloration in ecology and evolution, analysis of snakes and other elongate organisms lags behind taxa like insects and birds, specifically due to challenges in automating color quantification.

When using photography to collect color data, it is essential to identify which portions of a photograph are associated with the biological subject and which are calibration tools or part of the background (i.e., masking). Generally, standardizing preparation and photographing protocols reduces postural variation and enables comparison among specimens by facilitating the isolation of a biological subject. Morphological features, such as limbs, are often used as landmarks to identify color variation in homologous regions (Van Belleghem et al., 2018). Because snakes and many other animals lack limbs, their elongated body forms cannot be consistently positioned for photographic data collection. Snakes are usually coiled into circles or spirals for practicality, but the number, the diameter, and direction of the coils (clockwise or counterclockwise) vary greatly because snake length spans six orders of magnitude (Feldman et al., 2015). Such high disparity in morphology and posture hinders the application of traditional image processing techniques.

Recently, machine learning has facilitated the automated detection and visual categorization of biological information in large and complex datasets (Li et al., 2018). Machine learning can be performed by neural networks, which consist of processing nodes that distribute information to neighboring nodes, much like a human brain (Suk, 2017). These networks can be trained to perform specific tasks by providing a dataset in which the task has already been performed (i.e., a training set; see Section 6 for glossary). Then, the trained neural network can be applied to an unlabeled dataset to perform the same task (i.e., inference). By including sufficient variability in the training set, the trained neural network can robustly perform the task on diverse real-world biological data that vary in color, position, size, orientation, and resolution (Davis et al., 2020; Ditria et al., 2020; Kumar and Das, 2019).

Here, we present an automated and customizable workflow (`Batch-Mask`) using a region-based convolutional neural network (R-CNN) to identify and isolate pixels associated with biological specimens from photographs (Figure 1). First, we describe how we used labeled photographs to train the neural network for these diverse biological specimens (Section 3.2, Figure 2 A). Then, we use the inferred weights to automate masking of unlabeled photographs (Section 3.4, Figure 2 B). Finally, we demonstrate how `Batch-Mask` combines with other tools to automate biological image processing (Section 5, Figure 2 C). Due to their challenging variability, we use a diverse benchmark photographic dataset of 33 species of neotropical snake to assess automated masking and color pattern analysis (University of Michigan Museum of Zoology, Division of Herpetology et al., 2021). Because the weights are fine-tuned for a diverse dataset of coiled snakes, `Batch-Mask` is readily applied or trained to analyze other coiled forms, including whole organisms, organs, or tissues. By using a neural network to accommodate large amounts of variation in morphology and posture, this approach facilitates the automated analysis of highly diverse datasets for use in ecological and evolutionary analysis of color patterns.

**FIGURE 1.**
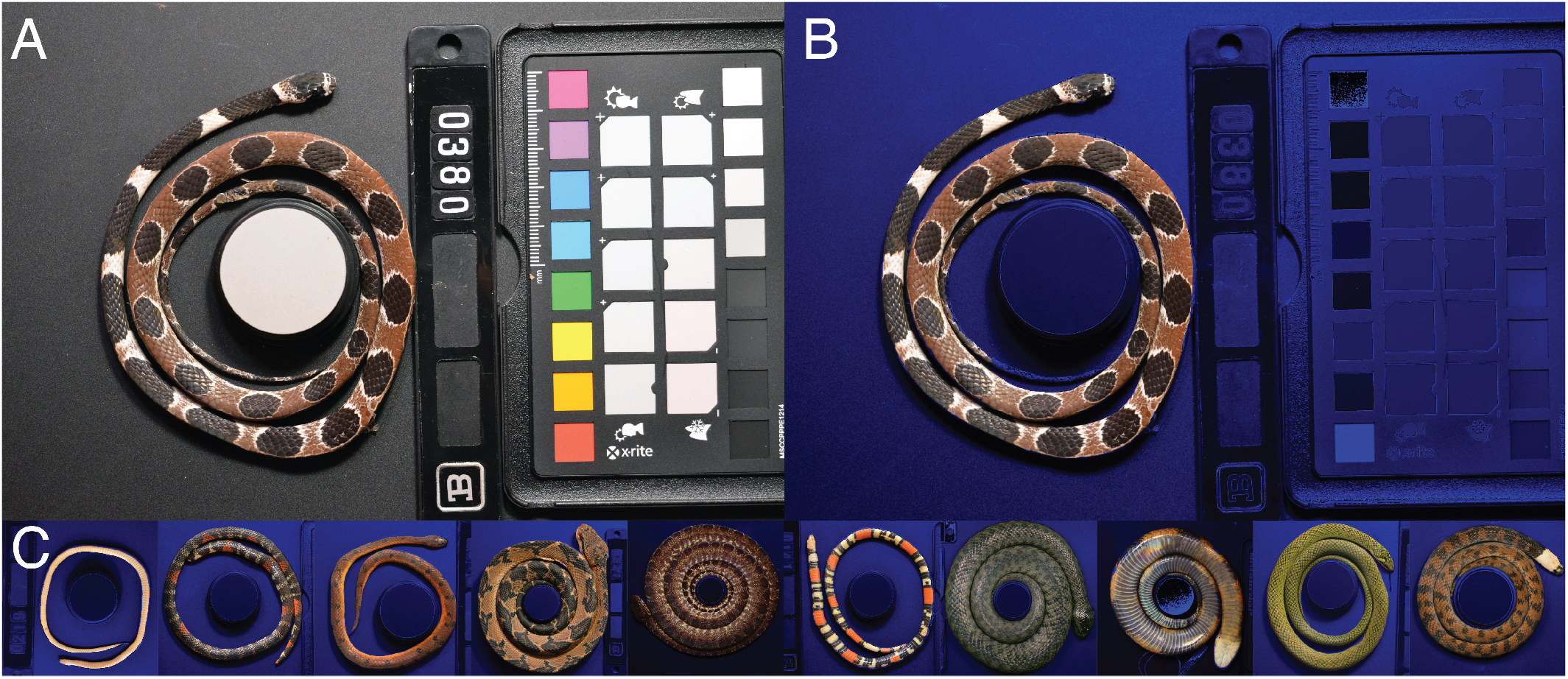
`Batch-Mask` fine-tuned weights use a neural network to take A) unlabeled photographs of circular or coiled biological specimens to generate B) a background-masked image. C) `Batch-Mask` is 60x faster than manual landmarking for specimens that vary in color, color pattern, thickness, coiling, and lighting conditions.

**FIGURE 2.**
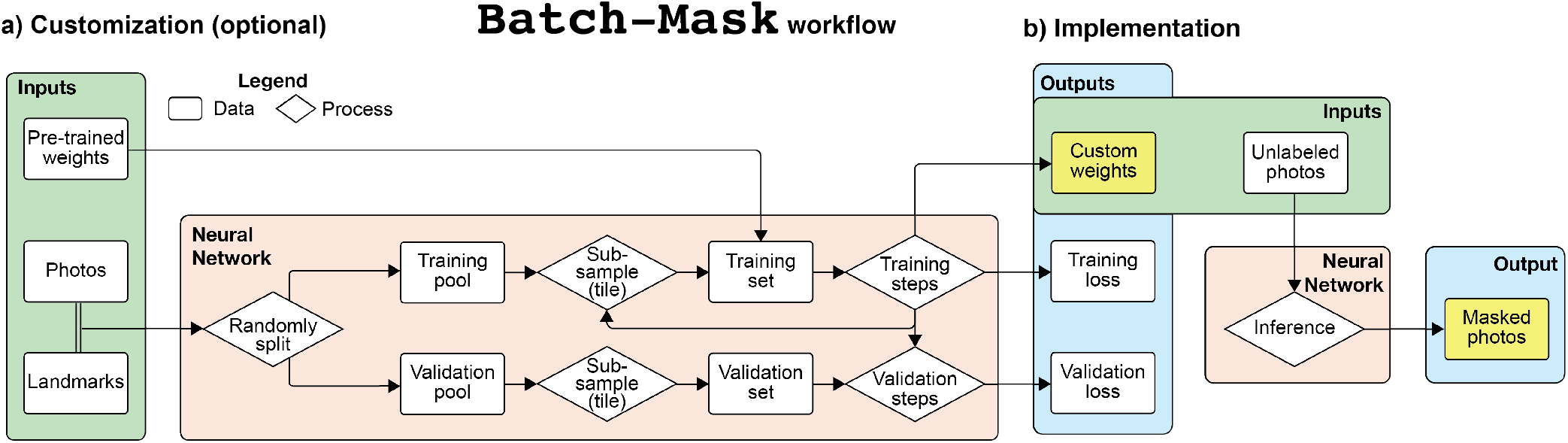
Summary diagram of the `Batch-Mask` work flow. A) Landmarked data from a few photographs are used to train a neural network and generate fine-tuned weights. This is unnecessary if the data set is visually similar to coiled snakes. B) Biological subjects are automatically isolated from an unlimited number of photographs without landmarks. Yellow boxes indicate outputs used for downstream processes

## 2 REQUIREMENTS AND INPUTS

### 2.1 Software

The `Batch-Mask` source code can be run through online cloud-based tools (such as Google Colab) to minimize hardware-induced performance constraints. Programs such as `ImageJ` (Schindelin et al., 2012) or `TPSDIG` (Rohlf) can be used to landmark the borders of a snake in training set photographs. Alternatively, the source code can be modified or run on a local machine, though we recommend access to a GPU. A list of required Python libraries is included in the source code). Any color calibration or color pattern analysis software can be used for downstream processing; here, we incorporated the `micaToolbox` (Troscianko and Stevens, 2015) plugin for `ImageJ` into a fully automated Python workflow.

### 2.2 Obtaining images

We trained and tested the neural network of `Batch-Mask` on an open-source dataset of Neotropical snakes (University of Michigan Museum of Zoology, Division of Herpetology et al., 2021). All specimens were photographed before preservation using a Nikon D7000 digital SLR camera (Nikon Inc., Melville, NY, USA) with a Coastal Optics UV-VIS-IR 60 mm F/4 macro lens (Jenoptik Optical Systems, Jupiter, FL, USA) using variable shutter speeds, F-stops, and ISO. Each specimen was illuminated from multiple angles using fluorescent and UV light bulbs. Each photograph contained one specimen, a set of color standards (X-rite Colorchecker Passport Pro), and a circular gray standard (40% Spectralon Diffuse Reflectance Standard) that also functioned as an object of known diameter for size calibration. Photographs were saved as JPG files, but `Batch-Mask` is compatible with any image file type.

To facilitate accurate downstream color comparisons, we wrote a custom macro in Photoshop (Adobe Inc., San Jose, USA) that uses the AutoColor tool to calibrate the color in each photograph. We also used the `OpenCV` (Bradski, 2000) `GaussianBlur` function with a 5×5 pixel kernel size to pre-process the photographs for the neural network.

### 2.3 Creating labeled data

To train and implement our model, we labeled a set of 151 photographs (Set 1 in the dataset) that included species with diverse colors and patterns and both dorsal and ventral views of each snake. We recommend that the dataset include a wide range of color, size, shape, and pattern variation to maximize generalizability and minimize overfitting (see Section 3.3).

We used the `tpsDig` program (Rohlf) to manually place landmarks along each side of the snake’s body to indicate the pixels associated with the snake. We then used a custom script to convert the tps outline into a JSON file.

Note that the output masks are highly dependent on the labeled data. In our dataset, lateral scales visible from the ventral view of the snake were excluded from the ventral landmarking due to substantially different color patterning (Figure S1), resulting in output masks of ventral photographs that identify regions most relevant to biological analyses, rather than the edge of the snake’s body.

## 3 TRAINING AND IMPLEMENTATION

To facilitate customization for biological subjects that differ greatly from the visual appearance of snake, we describe how to train the neural network to a customized ground truth dataset. However, if coiled or circular subjects are being analyzed, refer to Section 3.4 to begin implementation.

### 3.1 Creating training and validation sets

We randomly divided the 151 landmarked JSON files into a training pool of 135 labeled photographs and a validation pool of 16 labeled photographs, a ratio of 9:1 (see (Guyon, 1997) to determine optimal ratio).In every training step, we randomly selected a photograph from the training pool, then randomly sampled one 512 × 512 pixel square image (tile) from the photograph for the training set. We created one fixed validation set by randomly choosing 32 x,y coordinates and sampling a 512 × 512 pixel tile at each location from each photograph in the validation pool. Note that no validation tiles overlap with training tiles because they are sampled from different pools of photographs. However, validation tiles may stochastically overlap with each other.

### 3.2 Training the neural network

`Batch-Mask` utilizes a customized region-based convolutional neural network (R-CNN) model (He et al., 2017) to generate masks of snakes in photographs. This neural network uses the training process to fine-tune mask weights (*W*_*FT*_) from pre-trained weights (*W*_*PT*_) provided with Mask R-CNN (obtained from training on the COCO dataset (Abdulla, 2017)). On Google Colab (Abadi et al., 2015), we set the GPU count to 1 and the number of images per GPU to 1. Our learning rate was 0.0001 (see Section 4.1). All other parameters in the configuration file were left to their default values.

The number of validation steps must be equal to the number of tiles in the validation set so that loss is calculated on the full validation set for every epoch. Mask-RCNN suggests using twice as many training steps as validation steps (Abdulla, 2017). The number of training and validation steps in an epoch does not affect model accuracy, but if training and validation loss values converge after a single epoch, decreasing the number of training steps will reveal the progression of loss values. Decreasing training steps should be accompanied by decreasing validation steps, such that a roughly 2:1 ratio is maintained. If the loss values take more than 12 hours to converge, the number of training steps can be increased. If both the training and validation loss plateau at non-zero values, Section 4.1 discusses how model settings can be adjusted to increase accuracy.

The training that resulted in the best masks used 450 training steps and 50 validation steps for each epoch. We trained for 20 epochs, each lasting 1.21 hours. The training and validation losses plateaued at 16 epochs, after which the validation losses began increasing (likely due to overfitting). The weight values at 16 epochs were used for inference. Our training process duration was 24.2 hours.

### 3.3 Loss and model accuracy

We calculated training and validation losses using the ratio of correctly labeled pixels divided by all labeled pixels (Table S1).This loss equation does not penalize pixels incorrectly identified. The training loss values inform the training process, but the validation loss does not. We assessed model accuracy by comparing loss values after each epoch to assess asymptotic curves (Figure 3), indicating diminishing increases in accuracy with additional training.

**FIGURE 3.**
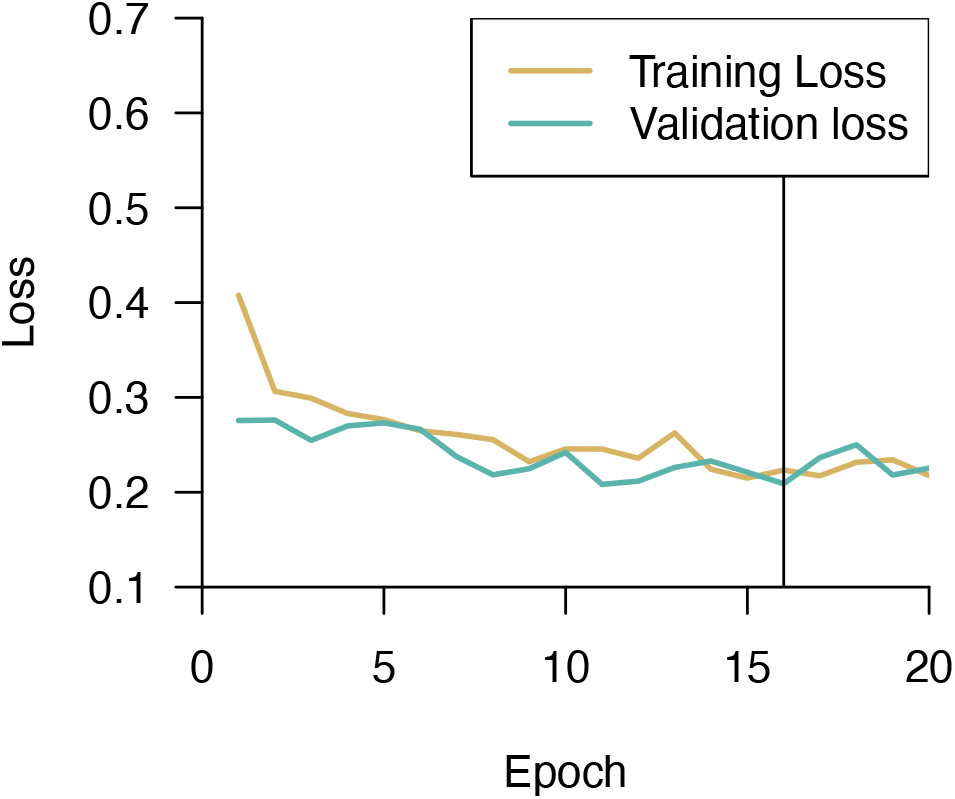
The loss calculations from a successful training process. The training loss is plotted in orange, the validation loss is plotted in teal. Epoch is indicated on the x-axis. Note that the training loss values decrease exponentially in the first epoch. The model weights corresponding to epoch 16, indicated by the black vertical line, were used for inference because this is the onset of the validation loss plateau.

Based on the validation loss, `Batch-Mask` was successful in rapidly isolating pixels associated with a biological subject from the background. To maximize the size of our training and validation sets, we used the validation loss at the epoch used for inference to represent the accuracy of the trained neural network, instead of inferring masks for a labeled test set. We qualitatively examined the output masks for an unlabeled test set (Section 3.4), which we found to be usable.

### 3.4 Implementation on Unlabeled Data (Inference)

To demonstrate the utility of our automated workflow to accurately process images outside of our training and validation sets, we implemented `Batch-Mask` on a test set of 50 unlabeled photographs (Set 2) (University of Michigan Museum of Zoology, Division of Herpetology et al., 2021), each subdivided into 212 tiles of 512 × 512 pixel resolution with 100 pixels of overlap with neighboring tiles in each direction. The implementation of the trained model required approximately 25 minutes to mask 50 unlabeled images. By comparison, generating a JSON file of ROIs for an equal number of images would require approximately 25 hours for a trained human to do by hand (based on landmarking rates in the training set).

Because there are no landmarks associated with these photographs, we created a Python workflow that displays a random subset of masks overlaid on original photos to qualitatively assess mask accuracy.

## 4 PARAMETER OPTIMIZATION

To assist with troubleshooting and customization of the workflow, we discuss settings as they relate to loss, mask outputs, and computation time. However, we note that this is not an exhaustive guide and other settings could also be modified (such as learning momentum, relative loss values, and mask shape).

### 4.1 Troubleshooting and modifying settings

The three parameters that have the most effect on output are learning rate, tile size (resolution), and tile overlap. earning rate controls the magnitude of the correction to the weights in response to a mismatch between training output and the labeled data. High learning rate values cause the training and validation loss values to diverge or wildly oscillate, while small values result in slower convergence but less oscillation. We recommend starting with smaller learning rate values and slowly increasing between training sessions to improve performance. Note, the Mask R-CNN code includes by default a learning rate decay throughout a training session, which was not modified for this method.

The resolution of each tile, the number of subdivisions, and the overlap between neighboring tiles sampled from each photograph are all interdependent. If the entire ROI is not identified (Figure 4 A, top) or if undesired regions are included, we recommend decreasing tile size (increasing the number of subdivisions for the same amount of overlap) to increase mask resolution. A higher resolution typically produces a more accurate model but requires exponentially more memory (Figure 4 A, bottom).

**FIGURE 4.**
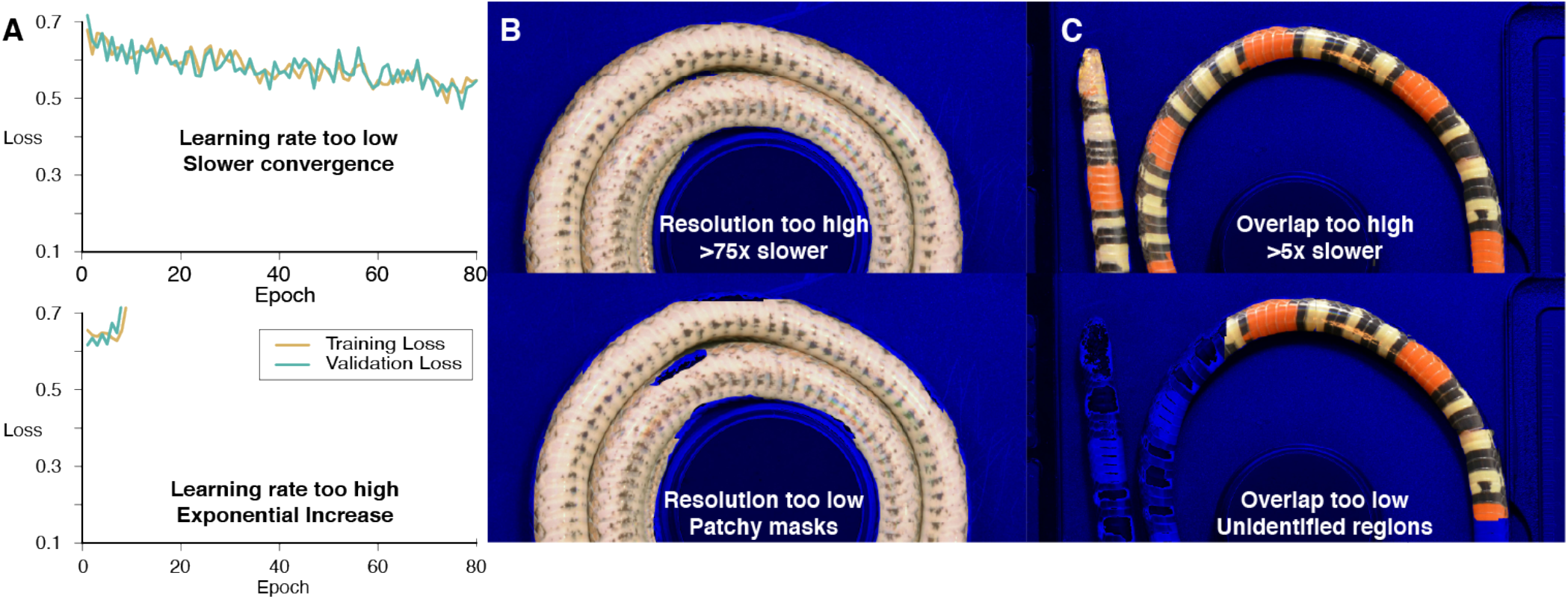
Visual guide to troubleshooting the training process. Each column in this diagram represents the indicators for tuning a different parameter. Loss plots can be used to troubleshoot a) Learning rate. Mask quality can be used to troubleshoot b) Subdivision/Resolution, c) Overlap. The top row displays the output indicating that the parameter should be increased. The bottom row displays the output indicating that the parameter should be decreased. Note that the loss plots are exaggerated to show the most recognizable patterns and were not generated by training results.

If mask inaccuracies correspond to the edges of subdivided image tiles, we recommend increasing overlap (the number of pixels shared between neighboring tiles). With higher overlap, features are viewed in multiple contexts, providing more opportunities for proper identification. Alternatively, decreasing overlap reduces computation time for acceptable output masks.

### 4.2 Overfitting

Accuracy is highest when the distribution of variation in the training and validation sets match. Overfitting occurs when the model is complex enough to memorize the entire training dataset, resulting in poor generalization (training loss decreases, but validation loss plateaus or increases after a plateau). To avoid overfitting, expand the size and variation within the training set (specimen size, shape, pattern diversity, color) to more accurately reflect the variation in the validation set. If diversity cannot be expanded with additional images, randomly changing the brightness and the hue of each image enhances useful variation in the training set.

### 4.3 Post-Processing

Because the loss calculation does not penalize non-ROI pixels misidentified as ROI pixels, the resulting inferred masks will likely include pixels outside of the snake. To eliminate these outliers, our script uses the `OpenCV` function `findContours` to identify the largest contiguous unmasked area and eliminate unconnected areas. This is helpful if portions of the background are being recognized incorrectly as ROI. If the ROI is in more than one contiguous piece, this function can be changed to recognize two or more unmasked areas.

## 5 DOWNSTREAM ANALYSES AND COMPATIBILITIES

Here, we demonstrate the ease of incorporating our `Batch-Mask` approach into a Python workflow for processing large datasets by analyzing the color pattern energy of snakes in the dataset (Figure 2 C).

Prior to analysis, we identified the gray size standard in each photograph using the Circle Hough Transform algorithm (available from `OpenCV` (Bradski, 2000)). The gray standard and snake ROIs were combined and exported into a single JSON file.

### 5.1 An automated workflow for color analysis

We incorporated the `micaToolbox` functions into the `Batch-Mask` workflow to combine color channels, scale the image by size, and create a single MSPEC file for pattern energy analysis of complexity in the snake ROI. To automate the process of generating MSPEC images, we modified the `micaToolbox` script to load the ROIs directly from the JSON file and batch-generate MSPEC files for multiple specimens at once.

To test whether the errors in machine-learned inference reduced the quality of color analyses, we compared hand-labeled masks to those produced by `Batch-Mask` for the photographs in the training set. Pattern energy as a function of pattern size for each color channel showed no significant differences between the hand-labeled and inferred datasets (paired t-tests, all *p >* 0.05, Figure 5). These results demonstrate that small pixel-wise differences between the hand-labeled and inferred datasets do not compromise the quality of downstream color analyses.

**FIGURE 5.**
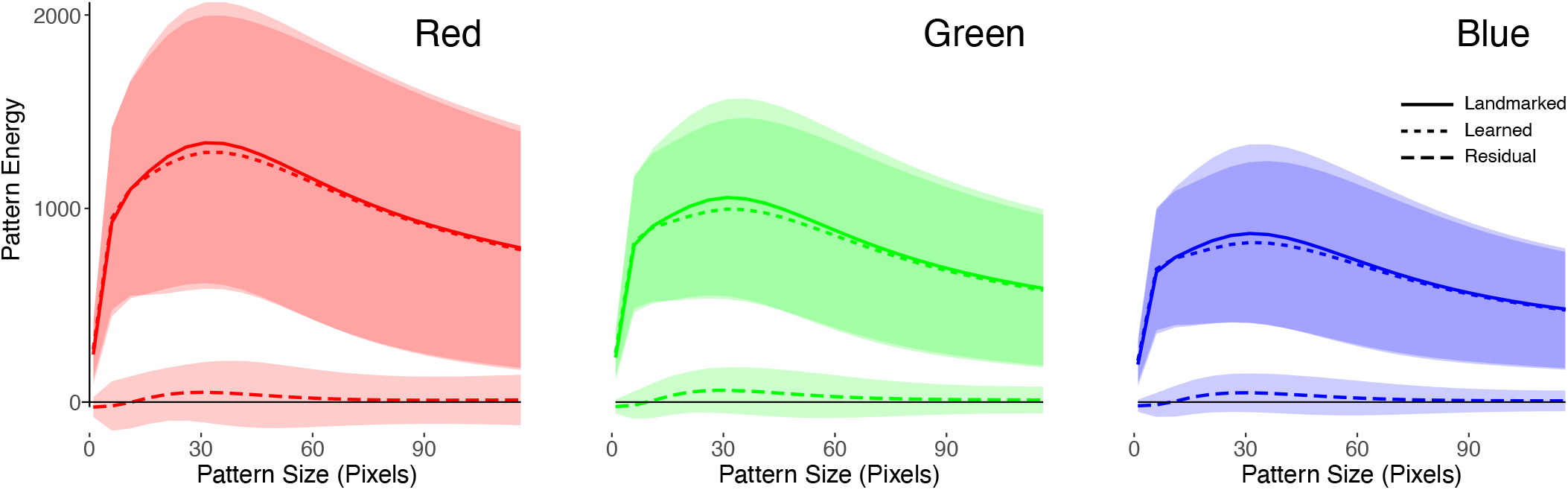
The mean residual differences (dashed line) between the pattern energies computed using the hand-labeled (solid line) and the inferred (dotted line) masks for the each photograph in the training dataset. Differences are sorted by color channel: red, green, and blue. The clouds represent the standard deviation from the mean for each pattern size in the color channel.

## 6 GLOSSARY

- **Mask R-CNN**: A Region-based Convolutional Neural Network that uses a mix of convolutional layers and fully connected layers to classify images.
- **Mask**: Binary array with the same dimensions as the image, with 1 indicating snake pixels and 0 indicated non-snake pixels.
- **Region of interest (ROI)**: Parts of an image outlined by a polygon or designated by a binary array.
- **Label**: Human-generated ROIs for both the training and validation sets.
- **Landmarking**: Identifying the locations of comparable morphological features among distinct biological specimens.
- **JSON**: JavaScript Object Notation formatted file comprising landmarking and image information.
- **Weights**: Values applied and changed during training to fit the neural network to data. Pre-trained weights (*W*_*PT*_) were provided with the standard Mask R-CNN model trained on the COCO dataset. Fine-tuned weights (*W*_*FT*_) fit during training are utilized, but not changed, during inference.
- **Training pool**: Set of labeled photographs from which sample tiles are extracted to generate the training set. There is no overlap between training and validation pools.
- **Training set**: Set of tiles sampled from photographs in the training pool. There is no overlap between training and validation sets.
- **Training step**: The neural network predicts the label for a tile from the training set, then updates the model weights if the model fails to match the true mask.
- **Validation pool**: Set of labeled photographs from which sample tiles are extracted to generate the validation set.There is no overlap between training and validation pools.
- **Validation set**: Set of labeled tiles used for the validation steps of the training process. There is no overlap between training and validation sets.
- **Validation step**: The neural network predicts the label for an image from the validation set but does not change the model weights. Validation steps calculate loss values on a labeled dataset that is distinct from the training set. This allows us to detect overfitting to the training dataset.
- **Loss**: A measure of the accuracy of model predictions for each pixel. Loss is calculated by the number of pixels shared by the label and the mask divided by the number of pixels in the label only, as in (He et al., 2017).
- **Epoch**: The interval of time after a certain number of training and validation steps have been completed. Model weights are saved after each epoch. Changing the number of training and validation steps per epoch changes how frequently model accuracy is assessed.
- **Training Process**: Using several epochs of training and validation to fine-tune model weights.
- **Inference Process**: Using fine-tuned weights to generate masks for data outside of the training and validation sets. Note that weights are not updated during inference.

## 7 ACKNOWLEDGEMENTS

We thank H. Crowell and J. Crowe-Riddell for helpful discussions, S. R. Manikandasriram and W. Weaver for comments on an early manuscript draft. This work was supported in part by a University of Michigan Undergraduate Research Opportunity Program and an MCubed grant to ARDR and TYM.

## 8 AUTHORS’ CONTRIBUTIONS

TYM and ARDR conceived the study; TR and TYM designed the methodology; TR, JDC, and TYM performed the analysis; JDC, TR, and TYM led manuscript writing. All authors contributed critically to the drafts and gave final approval for publication.

## 9 DATA AVAILABILITY

The masks associated with the photographs in the dataset are posted on Deep Blue Data https://doi.org/10.7302/3xwv-7n71. Code is posted on https://github.com/EMBiRLab/batch-mask.

