## Supplementary Material for "Batch-Mask: An automated Mask R-CNN workflow to isolate non-standard biological specimens for color pattern analysis"

### S1 | SUPPLEMENTARY MATERIAL

#### S1.1 | Dorsal and Ventral variability

The accuracy for the ventral photographs was lower than for dorsal photographs, likely due to variation in the original posture and lateral scale overlap in the ventral view (Table S1, Figure S1). In general, the landmarks on the dorsal view of a snake aligned perfectly with the edge of the snake's body, as there is typically no noticeable distinction between dorsal and lateral scales. This discrepancy caused the model to more often misidentify lateral scales in the ventral photographs, in turn resulting in lower accuracy. This discrepancy demonstrates that model accuracy greatly relies on original landmarking, and model accuracy increases when target features have clear visual differences from the background.

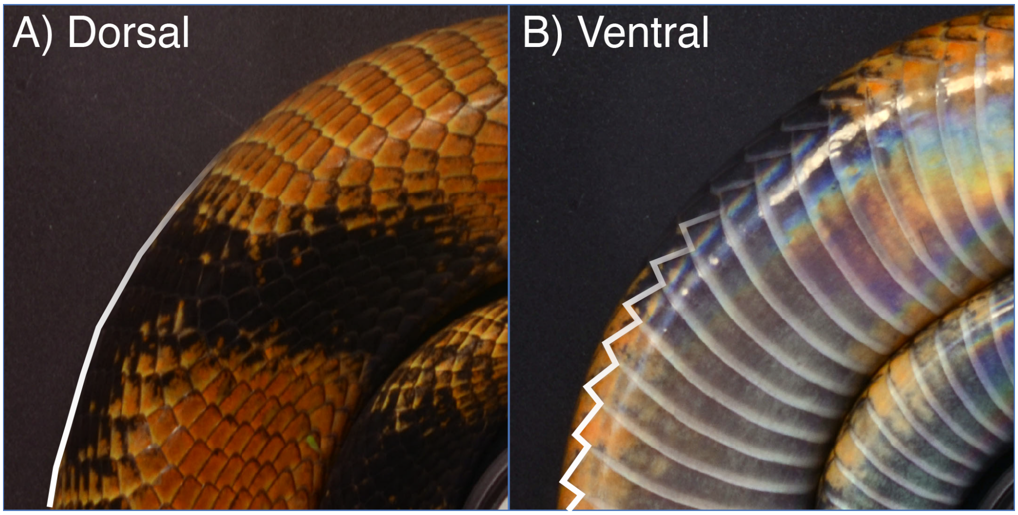


**FIGURE S1**Comparison between A) dorsal photograph of a snake and B) ventral photograph of the same snake. White outlines show landmarking borders. Note that ventral and lateral scales are less visually distinct from each other than the background is from most portions of the snake.

| Training set | Validation set | I/U | I/L |
|---|---|---|---|
| Dorsal | Dorsal | 88.1% | 96.0% |
| | Ventral | 74.1% | 93.9% |
| | Dorsal + Ventral | 80.2% | 94.8% |
| Ventral | Dorsal | 40.8% | 43.1% |
| | Ventral | 73.0% | 81.5% |
| | Dorsal + Ventral | 58.9% | 64.7% |
| Dorsal + Ventral | Dorsal | 87.6% | 95.3% |
| | Ventral | 80.8% | 91.2% |
| | Dorsal + Ventral | 83.4% | 93.0% |

**TABLE S1**Model accuracy using different combinations of training and validation images. Dorsal and ventral photographs were used to create different datasets (see Figure S1). Accuracy is calculated in two ways. Intersection/Union (I/U) by the number of shared pixels between the labeled data and the identified ROI (Intersection) divided by the total area identified by both the labels and the ROI (Union), which accounts for both Type I and Type II error. Intersection/Labeled (I/L) is calculated by intersection over Labeled pixels (equivalent to the accuracy metric in He et al. (2017)), which determines how many of the labeled pixels were accurately inferred, but does not penalize pixels incorrectly identified as ROI. The I/L value is used to train the model. Variability in the ventral landmarked data contributed to lower accuracy when these data were incorporated into the validation set. The identity of training and validation photos were consistent across experimental conditions. A mask trained on dorsal and ventral data was used for inference in Section 3.4.

**How to cite this article:** Curlis J.D. & Renney T.J., A.R. Davis Rabosky, and T.Y. Moore (????), `Batch-Mask`: An automated Mask R-CNN workflow to isolate non-standard biological specimens for color pattern analysis, *Methods in Ecology and Evolution*, *????;??:?–?.*
